# Development of White Matter Fiber Covariance Networks Supports Executive Function in Youth

**DOI:** 10.1101/2023.02.09.527696

**Authors:** Joëlle Bagautdinova, Josiane Bourque, Valerie J Sydnor, Matt Cieslak, Aaron F Alexander-Bloch, Max A Bertolero, Phil A Cook, Raquel C Gur, Ruben E Gur, Bart Larsen, Tyler M Moore, Hamsi Radhakrishnan, David R Roalf, Russel T Shinohara, Tinashe M Tapera, Chenying Zhao, Aristeidis Sotiras, Christos Davatzikos, Theodore D Satterthwaite

## Abstract

The white matter architecture of the human brain undergoes substantial development throughout childhood and adolescence, allowing for more efficient signaling between brain regions that support executive function. Increasingly, the field understands grey matter development as a spatially and temporally coordinated mechanism that follows hierarchically organized gradients of change. While white matter development also appears asynchronous, previous studies have largely relied on anatomical atlases to characterize white matter tracts, precluding a direct assessment of how white matter structure is spatially and temporally coordinated. Here, we leveraged advances in diffusion modeling and unsupervised machine learning to delineate white matter fiber covariance networks comprised of structurally similar areas of white matter in a cross-sectional sample of 939 youth aged 8-22 years. We then evaluated associations between fiber covariance network structural properties with both age and executive function using generalized additive models. The identified fiber covariance networks aligned with the known architecture of white matter while simultaneously capturing novel spatial patterns of coordinated maturation. Fiber covariance networks showed heterochronous increases in fiber density and cross section that generally followed hierarchically organized temporal patterns of cortical development, with the greatest increases in unimodal sensorimotor networks and the most prolonged increases in superior and anterior transmodal networks. Notably, we found that executive function was associated with structural features of limbic and association networks. Taken together, this study delineates data-driven patterns of white matter network development that support cognition and align with major axes of brain maturation.

## INTRODUCTION

Throughout childhood and adolescence, cerebral white matter expands dramatically and is extensively remodeled (1, 2). In tandem with this macrostructural expansion, microstructural properties of white matter also undergo substantial change, including increases in myelination, axonal density, and axonal caliber (3, 4). Cognition also develops rapidly during this period, with executive function undergoing a particularly protracted improvement throughout adolescence and young adulthood (5–7). The maturation of white matter architecture is thought to facilitate the efficient and coordinated relay of information between brain regions and to support the development of executive function (8). Quantifying how development of white matter structure supports executive function is a critical prerequisite for normative accounts of brain development as well as for studies of youth-onset psychiatric disorders that are characterized by both deficits in executive functions (9, 10) and differences within white matter (11).

Increasingly, we understand brain development as a spatially and temporally coordinated mechanism that progresses along major axes of brain organization (12–14). For example, cortical development has been shown to follow a sensorimotor-to-association (S-A) axis during childhood and adolescence, whereby lower-order sensory areas mature earliest and transmodal association areas show more prolonged age-related changes (12). Evidence for this hierarchical and heterochronous developmental pattern has now been observed using various measures derived from neuroimaging, including cortical thickness, functional connectivity, intracortical myelination, and intrinsic activity amplitude (12, 15, 16). Similarly, prior work on white matter development has also shown that different tracts exhibit asynchronous maturational timing, with projection and commissural tracts maturing earlier than association tracts (17). Although inconsistencies exist, available data suggest simultaneous inferior-to-superior and posterior-to-anterior patterns of white matter maturation (18). Notably, white matter maturation along these two anatomical axes could potentially reflect the maturation of grey matter along S-A axis (12). While findings thus point to a hierarchically organized developmental program, most prior work has relied on anatomical atlases to characterize white matter tracts, precluding a direct and data-driven assessment of how white matter development is spatially and temporally coordinated.

Understanding the ways in which white matter connections spatially covary is particularly important, given that covariance in brain structure is thought to result from coordinated maturational processes (19, 20). Furthermore, synchronized changes in grey matter structure and brain function are believed to be reciprocally influenced by the white matter bundles that directly connect to them (19). Therefore, characterizing the spatial and developmental covariance of white matter structure would provide a new perspective on white matter development that could not be intrinsically recovered based on tract-atlases only. Data-driven machine learning approaches are well-suited for this task, as they can accurately capture the covariance that exists across large-scale networks of structurally covarying areas. One recently developed approach for identifying covariance networks within high-dimensional neuroimaging data is non-negative matrix factorization (NMF), an unsupervised machine learning approach (21–23). NMF has been used to define data-driven covariance networks across the cortex using grey matter features such as cortical thickness and volume (24– 26), NMF has also been used to identify partitions of single anatomical regions (27–29). However, applications to white matter specifically remain quite rare (30).

Here, we leveraged NMF to uncover how structurally-covarying areas of white matter cooperatively develop to support executive function in a large sample of youth from the Philadelphia Neurodevelopmental Cohort (PNC). Specifically, we delineated white matter fiber covariance networks based on both macrostructural and microstructural fiber properties. The identified fiber covariance networks mapped well onto the known architecture of white matter while additionally capturing novel spatial and temporal patterns of coordinated maturation in structurally similar brain regions. We found that the largest developmental effects were concentrated in unimodal sensorimotor networks, while the most prolonged increases were located in superior and anterior transmodal fiber covariance networks. Finally, limbic and association fiber covariance networks were linked to individual differences in executive function. Together, our findings provide insight into how development unfolds across the brain’s white matter and contributes to the maturation of cognition in youth.

## RESULTS

We identified fiber covariance networks to study how structural properties of the brain’s white matter are related across spatially distributed areas and are refined in an age-dependent manner. To delineate these covariance networks, we applied the fixel-based analysis (FBA) pipeline (4) with single-shell three tissue constrained spherical deconvolution (31) to DWI data from 939 youth aged 8-22 years (see **Figure S1** for sample selection). This approach minimizes extra-axonal signal contributions from grey matter and cerebrospinal fluid, resulting in more accurate estimations of white matter structure for single shelled data. Moreover, unlike voxel-based diffusion modeling approaches, FBA gives a more accurate description of the underlying white matter geometry as it can identify multiple fiber populations within a voxel. These individual fiber populations in each voxel are referred to as “fixels” (4, 32). For each fixel, a fiber density and cross-section (FDC) measure was calculated that quantifies both microscopic (intra-axonal volume) and macroscopic (morphology differences in fiber bundle size) properties of white matter (4). We then applied orthogonal projective NMF (opNMF) (23), a data-driven unsupervised machine learning method, to fixel-wise FDC data to identify spatial networks where FDC covaries consistently across participants. The combination of a fixel-based approach with machine learning enabled us to both account for multiple crossing fibers within a single voxel and allow different fiber populations at a single voxel to be assigned to distinct covariance networks, resulting in an accurate representation of white matter maturation. opNMF produces a network matrix, containing the networks and their respective loadings on each fixel, and a participant-specific matrix, reflecting the average FDC value of each participant in a given covariance network (**Figure 1**). These participant-specific average FDC values for each network were then used as the dependent variable in subsequent group-level analyses.

**Figure 1.**
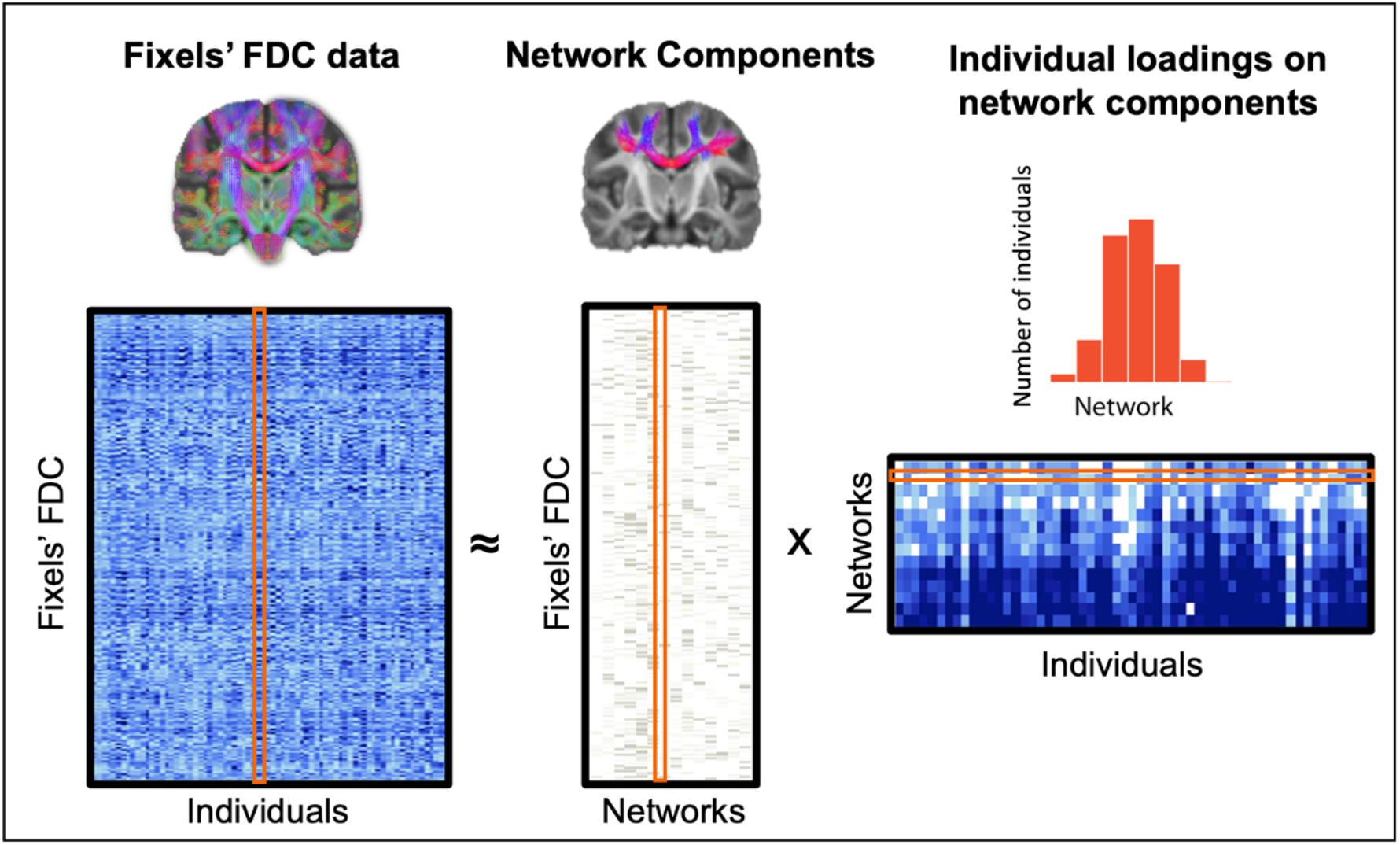
Identifying fiber covariance networks using non-negative matrix factorization. In this schematic, the original whole-brain FDC data for each fixel (rows) and for all individuals (columns) is fed into NMF, which then decomposes the data into a matrix of networks components and a matrix of individual loadings in each networks. The networks components matrix contains the loadings of each fixel on each of the 14 networks. Above the network components matrix is one example of fixel loadings onto a NMF network. The individual loadings matrix contains the participant-specific scores for each network. The histogram above shows a sample row of the matrix with scores for all participants in one network. Importantly, both output matrices are greater than or equal to 0; e.g., elements of the factorization are non-negative. The individual network loadings were used as the dependent variable in group-level analyses.

### NMF identifies anatomically meaningful fiber covariance networks

We applied NMF to segment whole-brain white matter into a low-dimensional number of networks, each comprised of white matter areas wherein FDC covaries in an organized fashion across individuals. Segmenting the brain’s white matter into 14 covariance networks parsimoniously captured FDC variability in a reproducible manner with low reconstruction error (see **Figure S2** for reconstruction errors of all network solutions). As in prior work using NMF on grey matter (21, 24), the 14 networks were sparse and highly symmetric bilaterally (**Figure 2**). Furthermore, they corresponded to known fiber bundles including commissural fibers (networks 1, 6 and 7), association fibers (5, 9, 10 and 14), projection fibers (3, 4 and 8), and cerebellar white matter (11, 12 and 13); one network included a mix of association and projection fibers (network 2). These covariance networks, however, also differed from known white matter bundle atlases in certain aspects. For example, different fiber bundles were in some cases aggregated into one network, such as the superior longitudinal and arcuate fascicules that form one covariance network (network 5). Moreover, in some components, the data-driven covariance networks also split different portions of a canonical fiber bundle into different networks, such as the inferior and superior/anterior cortico-spinal tract (networks 3 and 8). Thus, while covariance networks efficiently mapped major white matter bundles, they simultaneously revealed novel patterns of shared structural properties. To facilitate their identification in further sections, we labeled the 14 networks based on the tracts that are most strongly represented in them (**Figure 2**).

**Figure 2.**
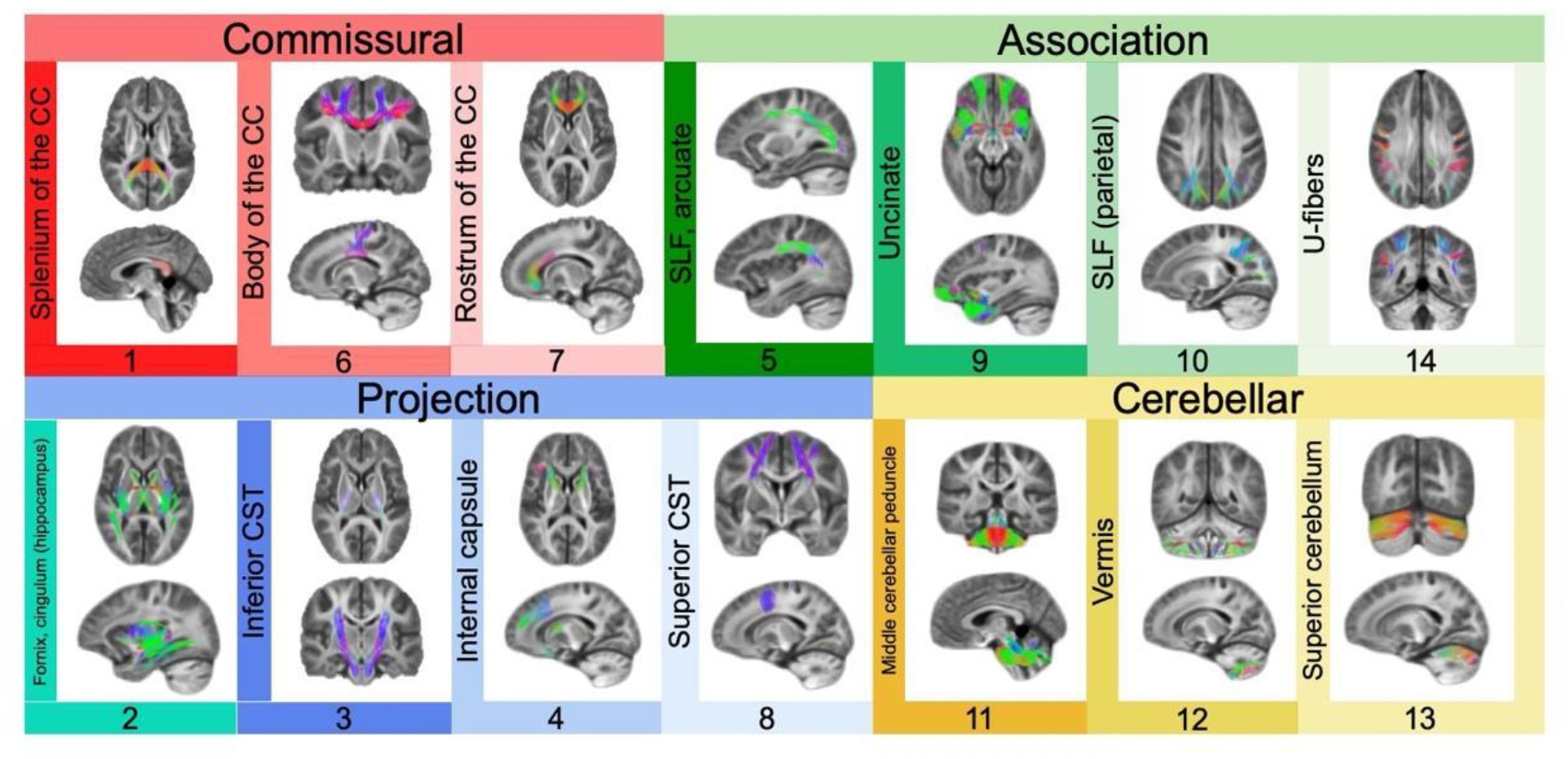
Delineating fiber covariance networks with orthogonal projective non-negative matrix factorization. opNMF yields a probabilistic parcellation such that each fixel receives a loading score onto each of the fourteen networks quantifying the extent to which the fixel belongs to each network. Here, the probabilistic parcellation was converted into discrete covariance network definitions for display by labeling each fixel according to its highest loading. The coloring of fixels is based on the Red-Green-Blue (RGB) convention which encodes the left-right, anterior-posterior, and inferior-superior directions, respectively. Networks identified include commissural bundles (1, 6, and 7), cerebellar white matter (11, 12, and 13), association bundles (5, 9, 10, 14, and 2), and projection bundles (2, 3, 4, and 8). Network 2 is included both in the association and projection networks because it encompasses the fornix and the cingulum (hippocampus). Network 10 refers to the parietal portion of the SLF.

### Fiber covariance networks show widespread development

A primary goal of this study was to characterize how macro- and micro-structural properties cooperatively develop across different white matter areas. Therefore, we investigated whether the identified fiber covariance networks displayed any developmental changes throughout childhood and adolescence. As brain maturation is a nonlinear process, we modeled age associations using generalized additive models (GAMs) with penalized splines, which rigorously capture nonlinear effects while avoiding over-fitting. All models included sex and in-scanner head motion as covariates. Our developmental models revealed that 12 out of 14 fiber covariance networks exhibited significant age-related changes in FDC (**Figure 3A** and **3B**), indicative of widespread white matter development. To determine the magnitude of the age effect in each covariance network, we computed effect sizes as the partial *R*^*2*^ – the proportion of variance explained by age. The largest effect sizes (partial *R*^*2*^ > 0.10) were found in networks encompassing the body of the corpus callosum, the superior longitudinal fasciculus and arcuate fasciculus, the splenium of the corpus callosum and the inferior corticospinal tract. These results indicate that most fiber covariance networks undergo maturational changes throughout adolescence, with the strongest age-related changes generally observed in networks connecting unimodal sensorimotor regions (with the exception of the network containing the superior longitudinal and arcuate fasciculi, which connects fronto-parietal regions).

**Figure 3.**
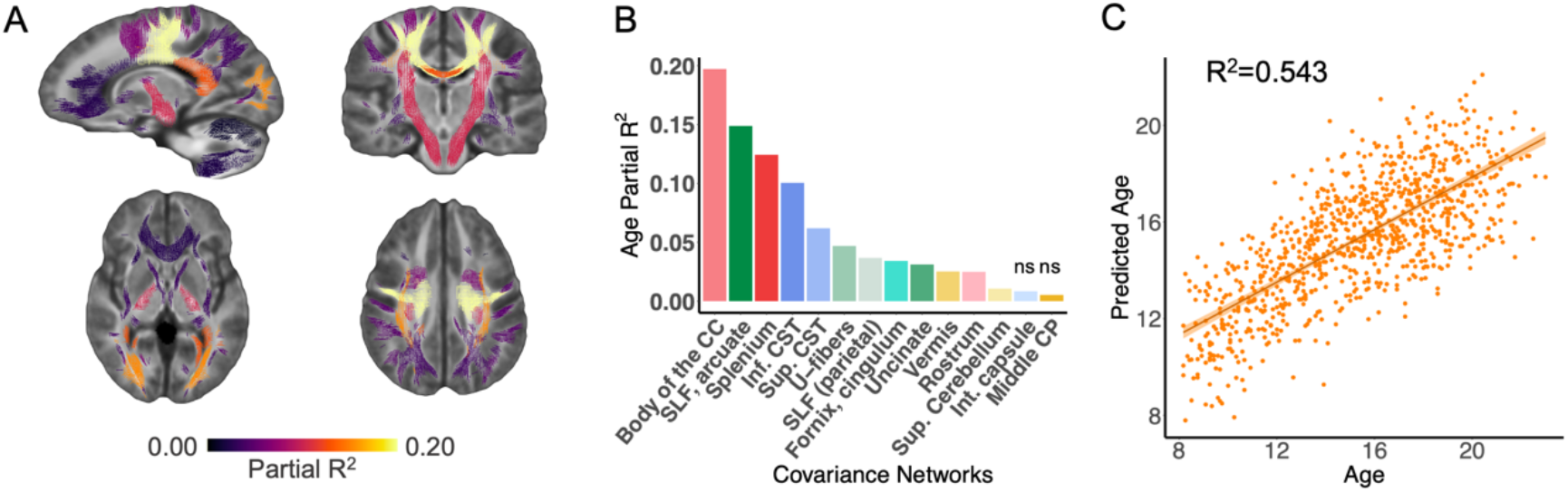
Developmental refinement of fiber covariance networks. **A)** Mass-univariate analyses using generalized additive models revealed that age was associated with significant changes in FDC in 12 out of 14 networks. The coloring of the covariance networks’ fixels is based on the variance explained (partial *R*^*2*^). Multiple comparisons were accounted for using the false discovery rate (*q*<0.05). **B)** Bar graph depicting the effect size (partial *R*^*2*^) of the developmental effect for each network. The greatest effect sizes were seen in networks such as the body of the corpus callosum, the superior longitudinal (SLF) and arcuate fasciculi, and the splenium of the corpus callosum (networks 6 and 5, and 1). Non-significant associations are marked by “ns”. **C)** We tested whether the multivariate signature of fiber covariance networks could predict age above and beyond sex and data quality by comparing a full model to a null model excluding the 14 covariance networks. We found a significant difference between a reduced covariate-only model (i.e., sex, motion, and image quality) and a full model that included both the fiber covariance networks and covariates *(F*=76.3, *df*=14, *p*<0.001). The proportion of variance in age explained by the 14 covariance networks was *R*^*2*^=0.543, resulting in a good correspondence between age and predicted age. Abbreviations: CC, corpus callosum; SLF, superior longitudinal fasciculus; CST, cortico-spinal tract; Sup, superior; Int, internal; CP, cerebellar peduncle.

The above quantified the magnitude of each individual fiber covariance network’s relationship with age. As a next step, we sought to determine the degree to which the multivariate signature of fiber covariance networks encoded development. We used a linear model to test whether participants’ covariance network scores, representing participants’ network-specific FDC, could predict age above and beyond demographic and diffusion data quality measures. We found a significant difference between a reduced covariate-only model (i.e., sex, motion, and image quality) and a full model that included both the fiber covariance networks and covariates *(F*=76.3, *df*=14, *p*<0.001). The proportion of variance in age explained by the 14 covariance networks was *R*^*2*^=0.543 (**Figure 3C**). These results demonstrate that fiber covariance networks effectively predicted participants’ age, and indicate that childhood and adolescence are a timeframe of robust refinement of fiber covariance network properties.

Having established that age effects were robust, we sought to obtain a more fine-grained representation of each covariance network’s maturational profile. We thus plotted network-specific developmental trajectories and quantified the direction and magnitude of maturational changes in FDC (**Figure 4**). The age-related changes were characterized by increasing FDC in most networks with the exception of the fornix/cingulum (hippocampus), vermis, and superior cerebellum where FDC decreased with age. To identify developmental windows of significant white matter maturation, we quantified the first derivative of the age smooth term, which represents the change in FDC at a given age. Age fit derivatives revealed distinct timings of developmental changes in different covariance networks. Specifically, the uncinate fasciculus, the parietal portion of the SLF, the splenium of the CC, the inferior CST, and superficial U-fibers showed increased FDC most significantly during childhood and the beginning of adolescence (i.e., from age 8 to prior to age 16). Other networks including the body of the CC, the SLF/arcuate fasciculus, the superior CST, and the rostrum of the CC displayed protracted increases in FDC throughout childhood and adolescence (i.e., until age 18). The FDC of the fornix/cingulum (hippocampus) network continued declining from age 8 through early adulthood (i.e., age 22). Conversely, the vermis and the superior cerebellum showed only brief windows of declining FDC. Thus, covariance networks captured temporally distinct patterns of coordinated white matter changes across different brain areas.

**Figure 4.**
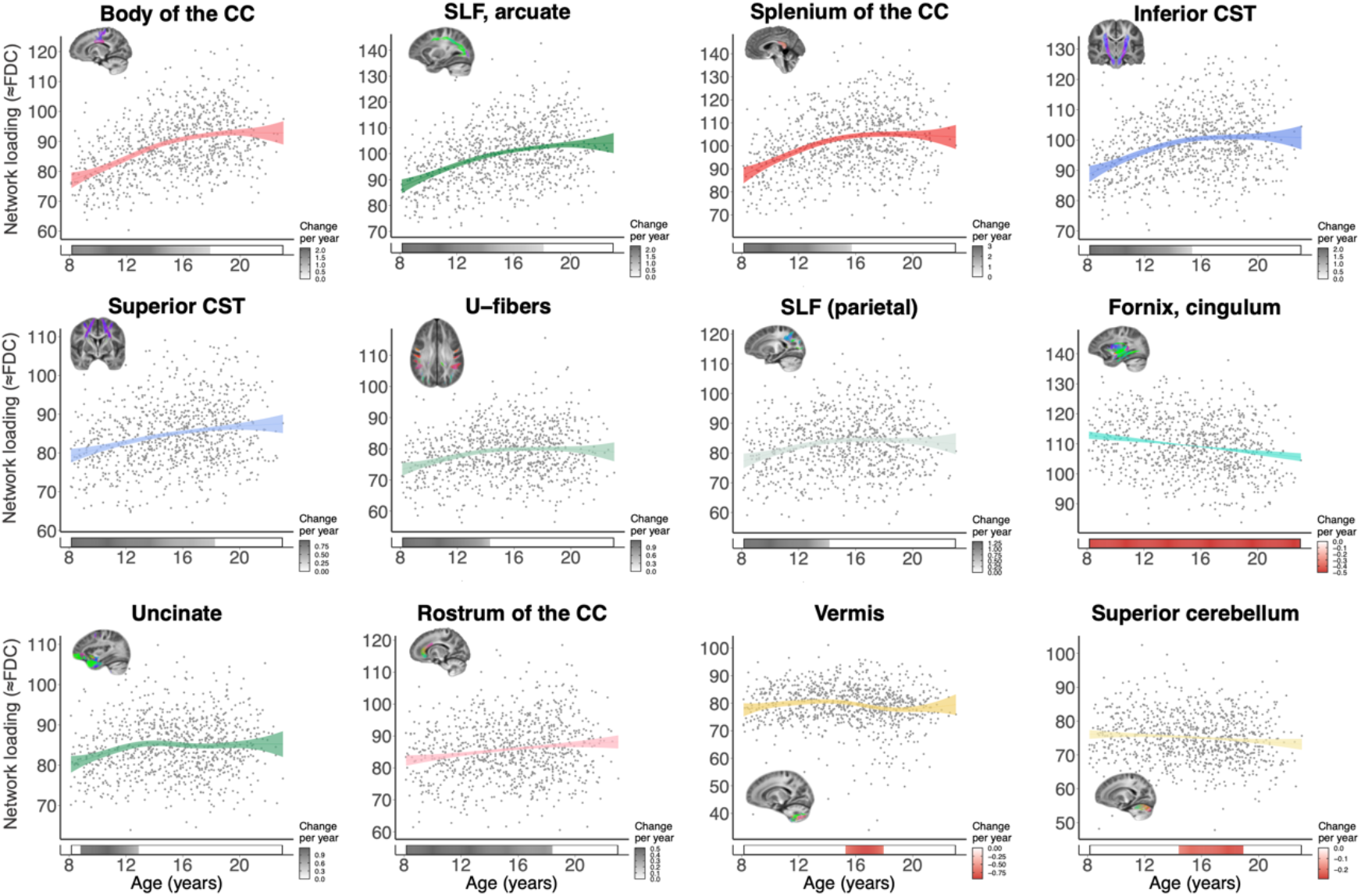
Development is associated with increased FDC in most fiber covariance networks. Plots display relationships quantified by generalized additive models between FDC and age for each covariance network. Significant age-related changes were characterized by increasing FDC in most networks, with the exception of the fornix/cingulum (network 2), vermis (network 12), and superior cerebellum (network 13). Bars below the x-axis depict the derivative of the fitted generalized additive model smooth function and correspond to developmental windows of significant white matter maturation. The filled portion of the bar indicates periods where the magnitude of the derivative is significant. A grey bar color indicates significant FDC increases (i.e., a positive derivative), and a red bar color indicates significant FDC decreases (negative derivative).

### Executive function is associated with variation in fiber covariance networks linking limbic and association cortex

Executive function is known to undergo considerable improvements during adolescence and young adulthood (5–7), and evidence from DTI studies suggests that these improvements are linked to white matter microstructural development (8). Based on these findings, we next examined whether variation in fiber covariance networks was related to differences in executive function. We found that better executive function was associated with higher FDC in 13 out of 14 covariance networks while controlling for age, sex, motion, and image quality (**Figure 5A** and **5B**). While significant, these univariate relationships showed modest effect sizes (partial *R*^*2*^ < 0.06). The three covariance networks with the greatest effect sizes include bundles such as the fornix/cingulum (hippocampus), the parietal section of the superior longitudinal fasciculus, and the combined superior longitudinal fasciculus and arcuate fasciculus network (**Figure 5D**). Thus, differences in executive function appear most strongly linked to limbic and association networks.

**Figure 5.**
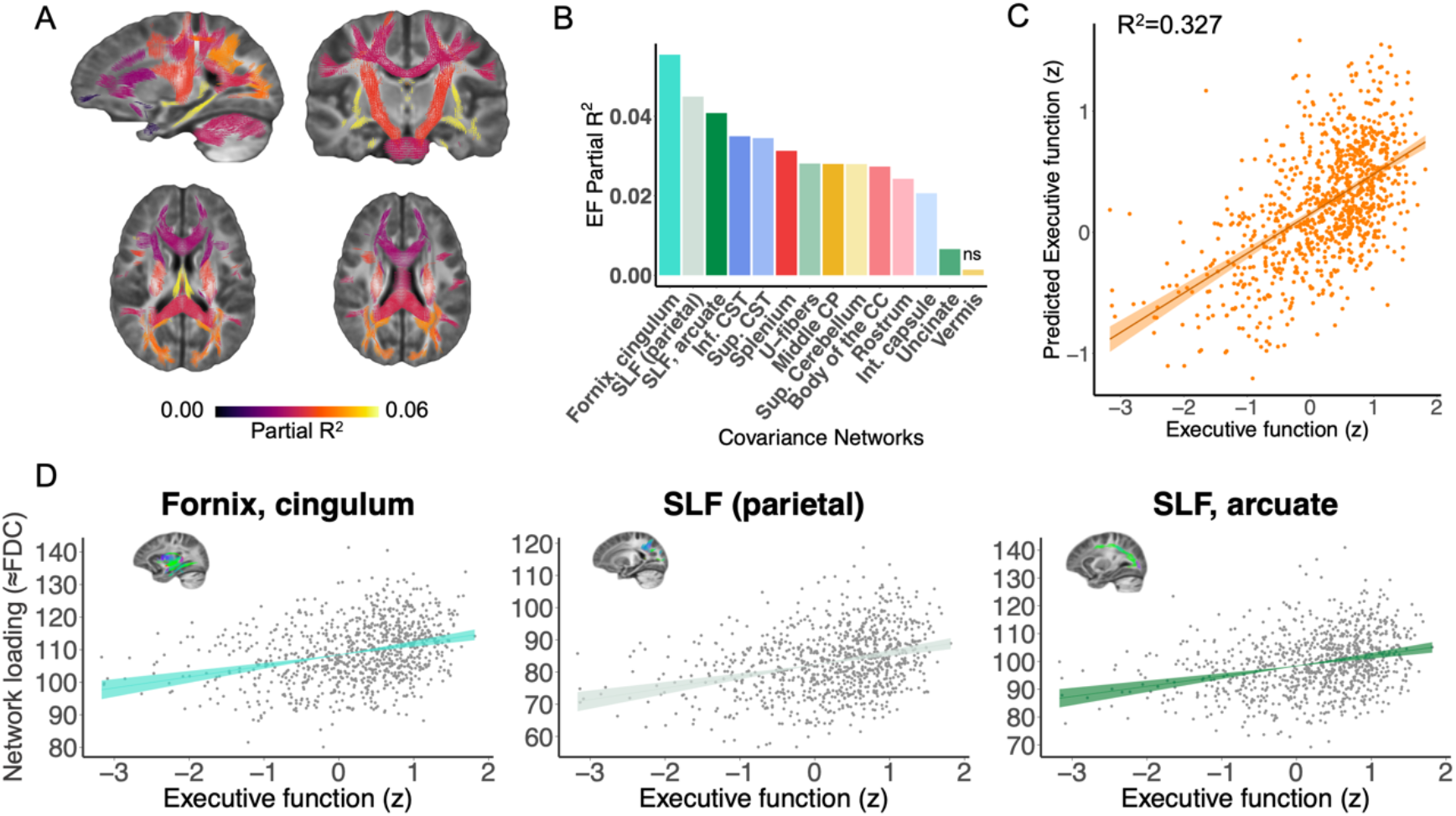
Fiber covariance network features are associated with executive function in youth. **A)** Univariate analyses using generalized additive models that controlled for sex, motion, and image quality revealed that executive function was associated with higher FDC in 13 out of 14 networks. The coloring of the covariance networks’ fixels is based on the partial *R*^*2*^ scores of executive function. Multiple comparisons were accounted for using the false discovery rate (*q*<0.05). **B)** Bar graph depicting the effect size of executive function for each network (partial *R*^*2*^). These partial *R*^*2*^ magnitudes were highest in association networks such as the fornix/cingulum (hippocampus), the parietal part of the superior longitudinal fasciculus, and the superior longitudinal and arcuate fasciculi (networks 2, 5, and 10). Non-significant associations are marked by “ns”. **C)** We tested whether the multivariate signature of fiber covariance networks could predict executive function above and beyond sex and data quality by comparing a full model to a null model excluding the 14 covariance networks. We found a significant difference between a reduced covariate-only model (i.e., sex, motion, and image quality) and a full model that included both the fiber covariance networks and covariates *(F*=6.56, *df*=14, *p*<0.001). The proportion of variance in executive function explained by the 14 covariance networks was *R*^*2*^=0.327, resulting in a good correspondence executive function and predicted executive function. **D)** The association between executive function and FDC is shown for the covariance networks with the highest partial *R*^*2*^ scores. Abbreviations: EF, executive function; CC, corpus callosum; SLF, superior longitudinal fasciculus; CST, cortico-spinal tract; Sup, superior; Int, internal; CP, cerebellar peduncle

Having examined associations between executive function and each individual fiber covariance network, we next evaluated whether the multivariate pattern of covariance network FDC could jointly predict executive function performance over and above age, sex, and data quality. We found that fiber covariance networks could explain over 30% of the variance in executive function beyond demographics and image quality (*R*^*2*^=0.327; *F*=6.56, *df*=14, *p*<0.001; see **Figure 5C**). Together, these findings suggest that fiber covariance networks may support executive function.

### Sensitivity analyses provide convergent results

As a final step, we conducted sensitivity analyses to evaluate potentially confounding variables. All associations between FDC and age remained significant following FDR correction after controlling for either total brain volume or maternal education (**Figure S3**). Similarly, the majority of the associations between FDC and executive function remained significant when total brain volume and maternal education were added as covariates (**Figure S4**). Exceptions included the rostrum of the corpus callosum, the internal capsule, the uncinate fasciculus and the U-fibers.

## DISCUSSION

We uncovered how structurally-covarying areas of white matter cooperatively develop during childhood and adolescence to support executive function. Specifically, we identified white matter structural networks based on covarying micro- and macrostructural-properties and delineated their age-dependent changes. The majority of these structural covariance networks showed age-related increases in FDC, which were further related to normative developmental improvements in youth executive function. Notably, the strongest developmental changes were mostly found in unimodal sensorimotor covariance networks, while the most prolonged developmental changes were found in superior and anterior transmodal networks. Furthermore, limbic and association networks were most strongly associated with executive function. As elaborated below, these findings suggest that white matter development follows inferior-to-superior and posterior-to-anterior axes that could influence the hierarchical maturation of the cortex.

### Networks derived from NMF complement anatomically defined white matter bundles

We derived data-driven fiber covariance networks using an advanced machine learning technique, opNMF (23). In comparison to the sparse, positively signed networks produced by opNMF, principal component analysis and other techniques produce widely dispersed networks with a mix of positive and negative weights, which often limit straightforward interpretations. Using a completely data-driven approach that did not incorporate explicit spatial constraints, opNMF revealed networks exhibiting high spatial contiguity within hemispheres and high bilateral symmetry. Our covariance networks aligned well with existing definitions of major white matter pathways, while also differing in certain aspects. First, since individual fiber populations were modeled within each voxel using fixel-based analysis, multiple fiber populations present within a given voxel could be assigned to distinct networks. This contrasts with most currently available white matter tract atlases, wherein each voxel is assigned to a single white matter bundle or network. Second, in our data-driven solution, several canonical fiber pathways were either aggregated into one network or had sections split across different networks. For example, the corticospinal tract was divided into an inferior and superior/anterior portion, and the superior longitudinal fasciculus divided into frontal and parietal connections – each with unique developmental trajectories.

Fiber covariance networks thus offer complementary information to standard anatomically defined atlases. Relative to anatomical atlases, data-driven atlases derived from covariance networks may more efficiently capture how brain areas function and mature together (33–35). Such a data-driven partition may be particularly useful to study brain organization from multi-modal neuroimaging-derived microstructural properties (36) and from comparison across different species (37). Future research could leverage our white matter network data-driven atlas to better understand how brain structural development underlies the development of functional brain networks.

### The maturation of fiber covariance networks follows principal axes of brain organization

We found that fiber covariance networks underwent different developmental patterns, with variations in both the magnitude and the timing of maturation. Generally, findings aligned with recent white matter development studies using advanced diffusion modeling strategies, which reported similar developmental timeframes (38, 39).

The greatest magnitudes of structural age-related change detected in the present study were found in networks primarily connecting unimodal sensorimotor cortices (i.e., body and splenium of the corpus callosum) and frontal to parietal regions (superior longitudinal and arcuate fasciculi). This aligns with age effects seen in functional networks, which were similarly largest in sensorimotor networks in the same cohort (16). Interestingly, from a spatial perspective, it appears that these networks are approximately located at intermediate stages along known inferior-to-superior and posterior-to-anterior axes of cortical organization. This is to be expected, given that the current study encompasses late childhood and adolescence, as opposed to early childhood (most inferior/posterior changes) and early adulthood (most superior/anterior changes). These major axes of brain organization have been highlighted from a lifespan developmental perspective of both gray and white matter (1, 40–42) and from specific developmental periods such as infancy (43), and youth (44, 45).

The timing of maturation varied considerably across fiber covariance networks, with some networks maturing in early childhood, some plateauing in adolescence, and some plateauing only by the end of adolescence. More specifically, fiber networks encompassing the cerebellum, fornix and parahippocampal cingulum showed either no change or decreasing FDC during adolescence. Conversely, some of the most prolonged age-related changes were observed in more superior and anterior fiber covariance networks linking to frontal and parietal areas (i.e., rostrum of the corpus callosum, superior longitudinal and arcuate fasciculi) as well as sensorimotor areas (body of the corpus callosum and superior corticospinal tract). Interestingly, the identified patterns of maturation across different covariance networks are again suggestive of a hierarchical patterning of development, whereby inferior and posterior white matter regions tend to mature earlier than superior and anterior areas. Of note, the S-A axis has been proposed as a more parsimonious way to capture the combination of inferior-to-superior and posterior-to-anterior developmental patterns (12). While further studies are needed to directly assess this, the data-driven networks identified in this study suggest that the timing of age-related changes may follow this hierarchical pattern, whereby white matter networks connecting to unimodal cortices mature earlier than networks connecting to transmodal cortices. Of note, it is likely that white matter connections linking to transmodal areas would show another wave of maturational changes going into the third decade of life (46). Further data-driven approaches on larger age ranges would therefore be optimally suited to capture this. Overall, these findings indicate that white matter structure is spatially and temporally coordinated in a manner that aligns with major inferior-to-superior and posterior-to-anterior axes of cortical organization. Results furthermore suggest that the sensorimotor-to-association pattern of development observed in the cortex may also exist within white matter, representing an exciting future area of investigation.

### Association covariance networks are linked to executive function

Having characterized the development of fiber covariance networks during adolescence, we lastly evaluated the cognitive impact of these developmental effects. While controlling for age, we found that an increase in FDC was associated with improved executive performance in all but one covariance network. The associations between FDC and executive function were most robust in fronto-parietal and temporal association networks, namely the parahippocampal cingulum as well as the superior longitudinal fasciculus and arcuate fasciculus. Higher FDC values may reflect increases in axonal packing and count and/or increases in the number of voxels that a bundle occupies (4). Of note, childhood developmental studies that have separately investigated FD, FC and their combination FDC, show more widespread and pronounced increases in cross-sectional fiber bundle size relative to increases in FD (47, 48). Applying this observation to the present results suggests that the changes in FDC may be mostly driven by changes in FC. Altogether, both microstructural (FD) and morphological changes (FC) may allow for more efficient signaling between distributed fronto-parietal and temporal regions that are critical for executive function. This result adds to prior work reporting associations between executive function and localized increases in white matter integrity in relatively small samples (8). Furthermore, it directly coheres with observations that greater cortical representation of association networks is linked to better general cognition in youth (49, 50) and suggests that the expansion of white matter underlying limbic and association cortex may contribute to these findings.

### Limitations

Despite the strengths of this study, several limitations should be noted. First, we used a cross-sectional design which precludes inference regarding within-individual developmental effects. However, it is worth mentioning that the present developmental effects of fiber density and cross-section agree with results from smaller-scale longitudinal studies (38, 47). Larger-scale longitudinal studies such as the IMAGEN study (51) and Adolescent Brain Cognitive Development Study (52) could eventually provide estimates of within-person change in fiber covariance networks at the population level. Second, the b-value of the diffusion imaging protocol is somewhat low (b = 1,000 s/mm^2^), which may result in less accurate FDC measures due to contamination from extra-axonal signal (38). In the present study, we addressed this caveat by using a single-shell multi-tissue modeling approach, which minimizes extra-axonal signal contributions. In addition, we chose to quantify white matter development using a combined measure of fiber density and fiber cross-section, which limited the biological specificity from the fiber density metric. Of note, prior work using fixel-based analysis has shown that the development of white matter is mostly driven by morphological changes from fiber cross-section (38, 47, 48).

### Summary

We introduced a data-driven approach for defining fiber covariance networks that are distinct from typical tract-based anatomical atlases and have unique associations with age in youth. Our findings suggest that developmental patterns of white matter follow large-scale anatomical gradients, in a way that may support heterochronous patterns of cortical maturation. Notably, specific limbic and association networks were associated with individual differences in executive function. Taken together, these results provide a novel framework to understand adolescent white matter organization and maturation using high-order diffusion modeling and data-driven multivariate analysis techniques.

## MATERIALS AND METHODS

### Participants

Among the original 1,601 individuals who participated in the Philadelphia Neurodevelopmental Cohort (PNC) (53), 340 were excluded due to clinical factors, including medical disorders that could affect brain function, current use of psychoactive medications, prior inpatient psychiatric hospitalizations, or an incidentally encountered structural brain abnormality. Among the 1,261 participants eligible for inclusion in this study, 174 participants were excluded for missing either a B0 field map or diffusion weighted MRI data. Data from the remaining 1,087 participants underwent both manual and automated quality assurance protocol for DWI (54, 55) and T1w datasets, which excluded 146 participants for poor quality data (see below for details and **Figure S1**). This set of exclusion criteria resulted in a final sample of 941 participants, with mean age 15.3 years, standard deviation (SD) = 3.4 years (n=522 females). All participants or their parent/guardian provided informed consent, and minors provided assent. All study procedures were approved by the Institutional Review Boards of both the University of Pennsylvania and the Children’s Hospital of Philadelphia.

### Cognitive assessment

The Penn computerized neurocognitive battery (CNB) was administered to all participants. The CNB consists of 14 tests adapted from tasks applied in functional neuroimaging to evaluate a broad range of cognitive domains (5, 56). These domains include executive control (abstraction and flexibility, attention, working memory), episodic memory (verbal, facial, spatial), complex cognition (verbal reasoning, nonverbal reasoning, spatial, processing), social cognition (emotion identification, emotion intensity differentiation, age differentiation), motor and sensorimotor speed. As in prior work (55, 57), the *z*-transformed accuracy and speed of each test were averaged to yield an integrated measure of cognitive efficiency (excluding motor and sensorimotor speed because they do not produce accuracy measures) that reflects accurate and rapid responding. These efficiency scores were then summarized by an exploratory factor analysis, which delineated four correlated factors (complex cognition, executive function, social cognition, and memory) (57). Two participants from the full n=941 sample had incomplete cognitive datasets, thus group-level statistical analyses examining associations between fiber covariance networks and executive function focused on the remaining 939 participants.

### Image acquisition

As previously described (53), all MRI scans were acquired on the same 3T Siemens Tim Trio scanner and 32-channel head coil at the Hospital of the University of Pennsylvania.

#### T1 weighted MRI

T1-weighted structural images were acquired prior to DWI acquisition with a 5-minute magnetization-prepared, rapid acquisition gradient-echo T1-weighted (MPRAGE) image with the following parameters: repetition time = 1810 ms, echo time = 3.51 ms, inversion time = 1100 ms, flip angle = 9 degrees, field of view = 180 × 240 mm, matrix = 192 × 256, slice number = 160, voxel resolution = 0.94 × 0.94 × 1 mm).

#### Diffusion MRI

Diffusion scans were acquired using a twice-refocused spin-echo (TRSE) single-shot echo-planar imaging (EPI) sequence (TR = 8100 ms, TE = 82 ms, FOV = 240 × 240 mm, matrix = 128 × 128, slices = 70, slice thickness/gap = 2/0 mm, flip angle = 90/180/180, voxel resolution = 1.875 × 1.875 × 2 mm, volumes = 71). A 64-direction set was divided into two independent 32-directions imaging runs - for a total scanning time of ∼11 minutes. Each 32-direction sub-set was chosen to be maximally independent such that they separately sampled the surface of a sphere. The complete sequence consisted of 64 directions with b = 1000 s/mm^2^ and 7 interspersed scans with b = 0 s/mm^2^.

#### Field map

In addition, a map of the main magnetic field (i.e., B0) using phase-difference images was derived from a double-echo, gradient-recalled echo (GRE) sequence, allowing us to estimate field distortions in each dataset (TR = 1000 ms; TE1 = 2.69 ms; TE2 = 5.27 ms; 44 slices; slice thickness/gap = 4/0 mm; FOV = 240 mm; effective voxel resolution = 3.8 × 3.8 × 4 mm).

### Image quality assurance

All T1-weighted anatomical images were independently rated by three highly trained image analysts as previously described in detail (58); participants with low quality structural images were excluded. Similarly, all dMRI images were subject to a rigorous manual quality assessment procedure involving visual inspection of all 71 gradient volumes (54). Each volume was evaluated for the presence of artifact, with the total number of volumes impacted summed over the series. Data was considered ‘‘Poor’’ if more than 14 (20%) volumes contained artifact, ‘‘Good’’ if it contained 1-14 volumes with artifact, and ‘‘Excellent’’ if no visible artifacts were detected in any volumes. All 941 participants included in the present study had diffusion datasets identified as ‘‘Good’’ or ‘‘Excellent’’ and also had less than 2 mm mean relative displacement between interspersed b = 0 volumes. Finally, to further mitigate potential effects of image quality on our DWI findings, measures of quality were included as covariates in all group-level analyses.

### Image processing

All images were processed with QSIprep, version 0.8.0RC3 (https://github.com/PennBBL/qsiprep) (59), which is based on Nipype 1.1.9 (60, 61). A total of 2 DWI series in the j-distortion group were concatenated, with preprocessing operations performed on individual DWI series before concatenation. Any image with a b-value less than 100 s/mm^2^ was treated as a b=0 image. MP-PCA denoising as implemented in MRtrix3’s *dwidenoise* (62) was applied with a 5-voxel window. After MP-PCA, Gibbs unringing was performed using Mrtrix3’s *mrdegibbs* (63). Following unringing, B1 field inhomogeneity was corrected using *dwibiascorrect* from Mrtrix3 with the N4 algorithm (64). After B1 bias correction, the mean intensity of the DWI series was adjusted so all the mean intensity of the b=0 images matched across each separate DWI scanning sequence.

Prelude from FSL (https://fsl.fmrib.ox.ac.uk/fsl/fslwiki/, version 6.0.3) was used to estimate the susceptibility distortion correction. FSL’s eddy was used for head motion and Eddy current correction (65). Eddy was configured with a *q*-space smoothing factor of 10, a total of 5 iterations, and 1000 voxels used to estimate hyperparameters. A linear first level model and a linear second level model were used to characterize Eddy current-related spatial distortion. *Q*-space coordinates were forcefully assigned to the shell. Field offset was attempted to be separated from participant movement. Shells were aligned post-eddy. Eddy’s outlier replacement was run (65). Data were grouped by slice, only including values from slices determined to contain at least 250 intracerebral voxels. Groups deviating by more than 4 standard deviations (SD) from the prediction had their data replaced with imputed values. Final interpolation was performed using the jac method.

The DWI time-series were resampled to the anterior-to-posterior commissure (ACPC) generating a preprocessed DWI run in AC-PC space with 1.25mm isotropic voxels. Several confounding time-series were calculated: framewise displacement (FD) based on the preprocessed DWI using the implementation in Nipype (following the definitions by (66)) and the number of bad slices in raw images. These two metrics - mean framewise displacement and number of bad slices - were included in all group-level statistical analyses.

### Fixel-based analysis

We pursued a modeling technique that models information from multiple distinct fiber populations within a given voxel: the fixel-based analysis (FBA) pipeline (4). In this pipeline, an individual fiber population within a voxel (fixel) is derived from fiber orientation distributions (FODs) estimated by a CSD technique (31). This approach minimizes extra-axonal signal contributions from grey matter and cerebrospinal fluid, resulting in more accurate estimations of white matter structure for single shelled data. Moreover, unlike voxel-based diffusion modeling approaches, FBA gives a more accurate description of the underlying white matter geometry as it can identify multiple fiber populations within a voxel.

Diffusion images were further processed using MRtrix3 (v3.0RC3, http://www.mrtrix.org/) (67) according to the FBA pipeline (4), which involves the following steps. First, we calculated study-specific response functions for white matter, gray matter, and cerebrospinal fluid (68) using data from 30 representative study participants (1 female and 1 male from each of fifteen age bins: 8-9 years old, 9-10 years old, up to 22-23 years old). Study-specific average response functions were then used to estimate FODs for all individuals with single-shell three tissue CSD (31). Following group-wise intensity normalization of the FOD images, we generated a study-specific FOD template (with the same 30 participants) and each individual’s FOD image was registered to the study template. The resulting participants’ transformed FOD images in template space were segmented along their main directions to delineate individual fiber bundle elements in each voxel, referred to as “fixels” (main manuscript **Figure 1**) (4, 32). Apparent fiber density (FD) was calculated for each fixel as the integral of the FOD lobe. Fiber cross-section (FC) was computed using the spatial warps generated during registration of each participant’s FOD image towards the common FOD template (4). Finally, a combined fiber density and cross-section (FDC) measure was calculated that quantifies both microscopic (intra-axonal volume) and macroscopic (morphology differences in fiber bundles) properties of white matter. FDC has previously been found to be more sensitive to detect change in white matter properties relative to FD and fiber cross-section alone (4). After calculating FDC for each fixel, the FDC values were smoothed to increase the signal-to-noise ratio. However, it is important to smooth fixel-specific metrics only with other fixels that share common streamlines, and not with all adjacent fixels (32). Therefore, to appropriately smooth fixels’ FDC values, we first generated a whole-brain probabilistic tractogram from the FOD template – in this case, a 20-million streamlines tractogram. The tractogram was reduced to a 2-million streamlines tractogram using the Spherical-deconvolution Informed Filtering of Tractograms algorithm (SIFT) (69) for which streamlines density is proportional to FD as estimated with CSD. Lastly, we computed a fixel-fixel connectivity matrix based on the reduced tractogram to inform smoothing of the fixel data at 10mm Full-Width at Half Maximum. Smoothed FDC metric for 602,229 fixels was used as input to the unsupervised machine learning approach using opNMF.

### Non-negative matrix factorization

We employed orthogonal projective NMF (23) to identify networks where fibers’ FDC covaried consistently across participants. opNMF produces sparse, positively-signed components that form a purely additive and non-overlapping parts-based representation (23). opNMF decomposes the input matrix X containing fixel-wise FDC estimates − of dimensions F x N (F = 602,229 fixels, N = 941 participants) − into a network matrix W (of dimension F x K; K = user-specified number of networks) and a weight matrix H (of dimension K x N; main manuscript **Figure 1**). The network and weight matrices are estimated such that their multiplication reconstructs the input matrix as best as possible by minimizing the reconstruction error between the original and the reconstructed input – the Frobenius norm. The network matrix W contains the estimated non-negative networks and their respective loadings on each of the fixels. This probabilistic (soft) definition of networks can be converted into discrete (hard) network definitions for visualization by labeling each fixel according to its highest network loading. The weight matrix H contains participant-specific scores for each network.

These participant-specific scores are equivalent to an average of FDC values for each covariance network. Consequently, the higher the participant score on a network, the higher that participant’s FDC value within that network. The participants’ scores from matrix H for each network were then used as the dependent variable in the subsequent group-level univariate analyses.

We used ConFixel (70) (https://github.com/PennLINC/ConFixel) to convert participants’ FDC data at every fixel into a readable Hierarchical Data Format 5 (HDF5) file format for NMF. Then, we used Brainparts (https://github.com/asotiras/brainparts) and Matlab R2018a to run the opNMF decomposition. Consistent with prior studies using this technique (21, 71), we evaluated multiple NMF solutions from 2 to 30 networks (in steps of two) to obtain a range of possible solutions. We selected the optimal network solution using the reconstruction error criterion. As previously detailed (21), this involves a visualization of the residual error after estimating different numbers of components (**Figure S2**). The inflection point of the reconstruction error slope indicates the optimal number of components, given that adding more components than the intrinsic data dimension only results in minor decreases in reconstruction error.

### Statistical analysis

After delineating fiber covariance networks, we evaluated associations between network FDC and both age and executive function. As brain maturation is a nonlinear process, we modeled age associations using generalized additive models (GAMs) with penalized splines in the R package “mgcv” (72). GAMs assess a penalty with increasing nonlinearity to avoid overfitting the data. Due to this penalty, GAMs designate nonlinearities only when they explain additional variance in the data above and beyond linear effects. All models used up to four basis functions, which were selected using the restricted maximum likelihood framework (REML) to produce estimates of variance parameters. For each white matter covariance network, we examined associations between age and FDC while controlling for sex, mean framewise displacement during the diffusion scan, and the number of slices with signal dropout (“bad slices”) observed in diffusion volumes, using the following formula:

NMF fiber covariance network score = spline(age, *k*=4) + sex + motion + bad slices + error.

To identify developmental windows of significant white matter maturation, we quantified the first derivative of the smooth age term, which represents the slope of the spline fit, at every age. Using the *gratia* package in R, we operationalized the window of significant age-related change as the period at which the 95% point-wise confidence interval of the spline’s estimated slope did not include 0.

To investigate whether executive function performance was associated with each network’s FDC above and beyond the effect of age, we included the executive efficiency factor score as a linear variable in the model above. In each set of analyses, multiple comparisons were controlled for using the False Discovery Rate (*q*<0.05) and effect sizes were quantified as the partial *R*^*2*^. The partial *R*^*2*^ is the proportion of variance explained by a full model that is not explained by a reduced model. For example, when investigating the relationship between age and FDC, the full GAM model includes age and the aforementioned covariates while the reduced model includes only covariates.

Following the investigation of the developmental and executive function effects of each covariance network separately, we next explored how well FDC of all these covariance networks jointly encode age and individual differences in executive function. Using a linear model, we tested whether the 14 covariance networks’ FDC jointly predicted age above and beyond sex, in-scanner motion, and quality of diffusion data. We used an F-test to compare the full linear model predicting age to a reduced model that excluded the 14 covariance networks. Similarly, we explored whether a network’s FDC predicted executive function above and beyond covariates and compared this full model to a reduced model excluding the 14 covariance networks. Finally, we calculated the proportion of variance explained (partial *R*^*2*^ coefficient) by the 14 covariance networks in predicting age and executive functioning.

### Sensitivity analyses

We conducted sensitivity analyses to ensure our results were not influenced by confounding variables. First, we repeated all group-level statistical analyses while including maternal education level as an additional covariate. Second, we included total brain volume in all group-level analyses to evaluate whether our results were driven by gross differences in brain volume.

## Code accessibility

To ensure reproducibility, the code for image preprocessing, evaluating NMF solutions, and all statistical analyses are available at https://github.com/PennLINC/Fixel_NMF_development. The Philadelphia Neurodevelopmental Cohort (PNC) (53) data are publicly available in the Database of Genotypes and Phenotypes (https://www.ncbi.nlm.nih.gov/projects/gap/cgi-bin/study.cgi?study_id=phs000607.v3.p2).

## ACKNOWLEDGMENTS

This study was supported by grants from the National Institutes of Health: R01MH112847, R01MH120482, R37MH125829, R01EB022573, R01MH113550, R01MH123550, RF1MH116920, K99MH127293. Additional support was provided by the AE Foundation, the Penn Center for Biomedical Image Computing and Analytics, and the Penn/CHOP Lifespan Brain Institute. JB was supported by a Canadian Institutes of Health Research Postdoctoral Fellowship CIHR-396349. VJS was supported by a National Science Foundation Graduate Research Fellowship DGE-1845298. DRR was supported by R01MH119185 and R01MH120174; AS was supported by R01 AG067103.

## SUPPLEMENTAL FIGURES

**Figure S1.**
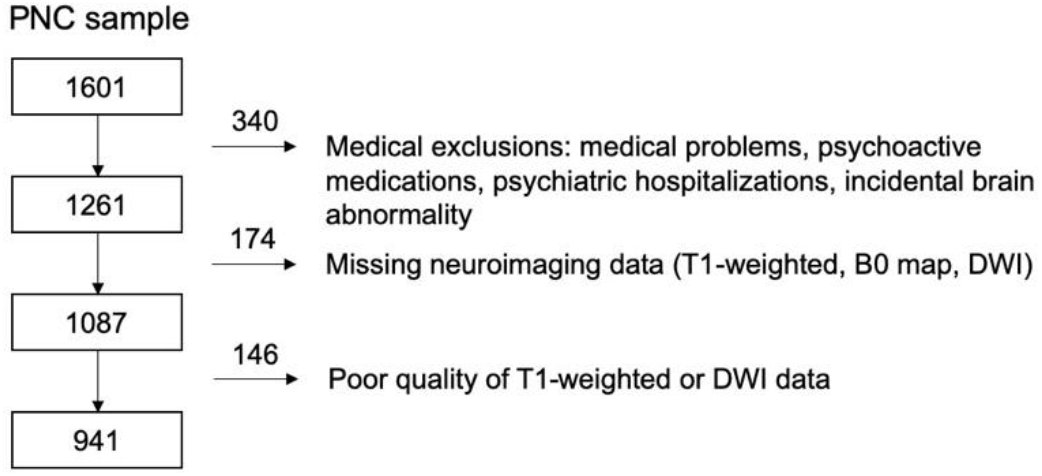
Sample construction. Among the original 1,601 participants from the PNC, 340 participants were excluded due to clinical factors, including medical disorders that could affect brain function, current use of psychoactive medications, prior inpatient psychiatric hospitalizations, or an incidentally encountered structural brain abnormality. Among the 1,261 participants eligible for inclusion, 174 participants were excluded for missing either a B0 field map, and/or diffusion images. The remaining 1,087 participants underwent a rigorous manual and automated quality assurance protocol for DWI datasets, which excluded 146 participants for poor data quality. This set of exclusion criteria resulted in a final sample of 941 participants.

**Figure S2.**
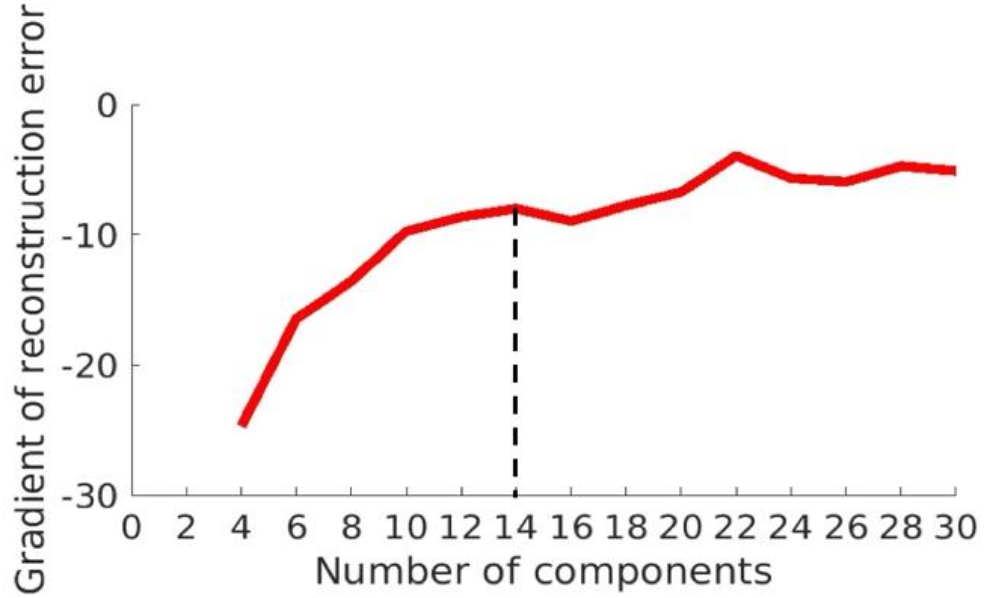
Gradient of reconstruction error for opNMF solutions. Reconstruction error is plotted for opNMF solutions ranging from two to thirty components. The gradient is the difference in reconstruction error of the X matrix (input data) as the opNMF solution increases by 2 components. The y-axis of the plot ranges from -100 to 0. However, to better visualize the differences in reconstruction errors between the different solutions, the y-axis was cropped to -30. As expected, reconstruction error plateaus as the number of components increases. The reconstruction error between the 10- to 30-components are fairly similar. We chose the 14-components solution as it is the most parsimonious solution before a small drop in reconstruction error. Accordingly, the 14-network solution was used for all subsequent analyses.

**Figure S3.**
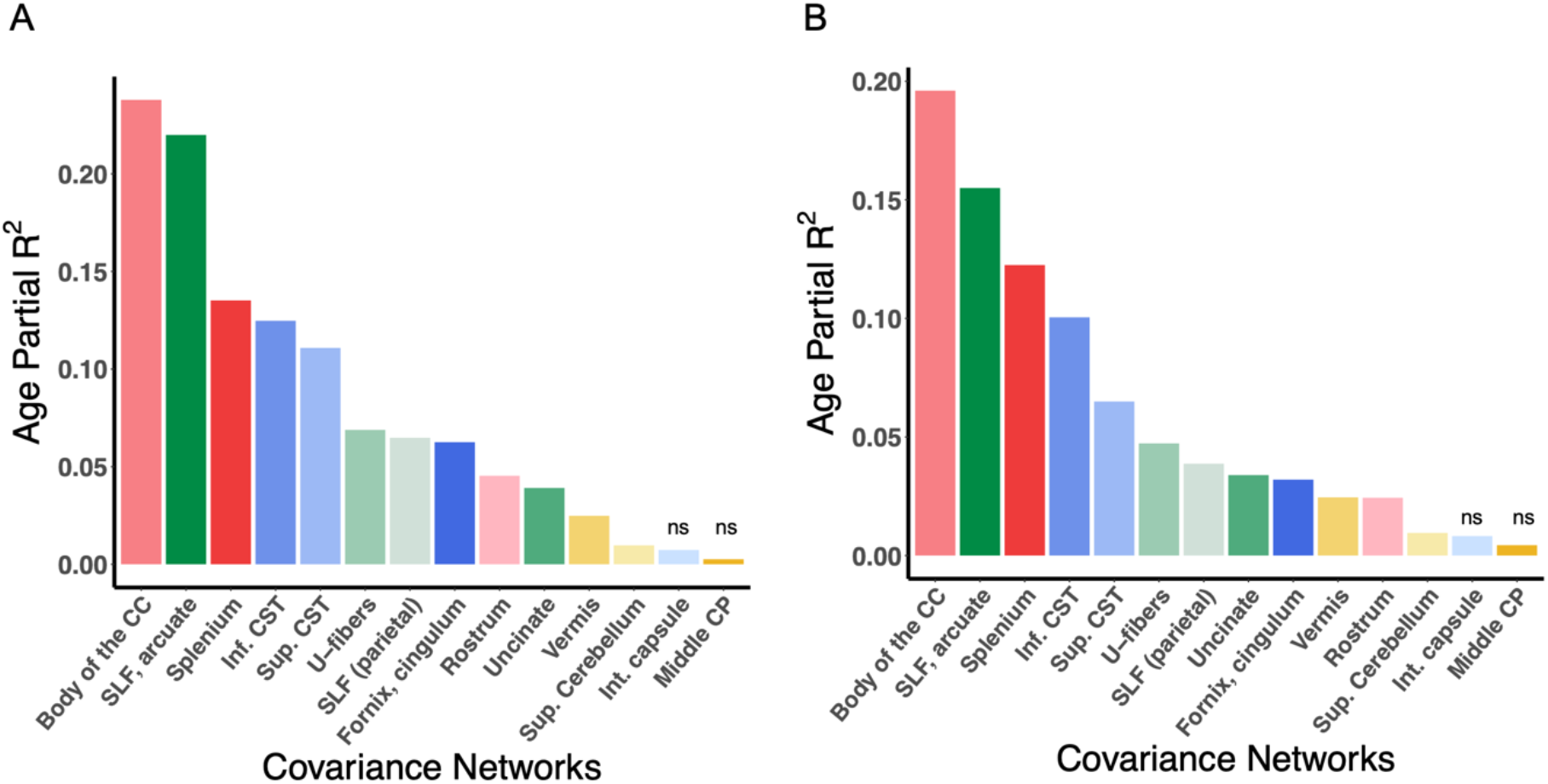
Effect sizes (partial *R*^*2*^) of age in each fiber covariance network controlling for total brain volume and maternal education. **A)** Bar graph depicting the effect size of the developmental effect for each network (partial *R*^*2*^) while controlling for total brain volume. **B)** Bar graph depicting the effect size of the developmental effect for each network (partial *R*^*2*^) while controlling for maternal education. Non-significant associations are marked by “ns”. Abbreviations: CC, corpus callosum; SLF, superior longitudinal fasciculus; CST, cortico-spinal tract; Sup, superior; Int, internal; CP, cerebellar peduncle.

**Figure S4.**
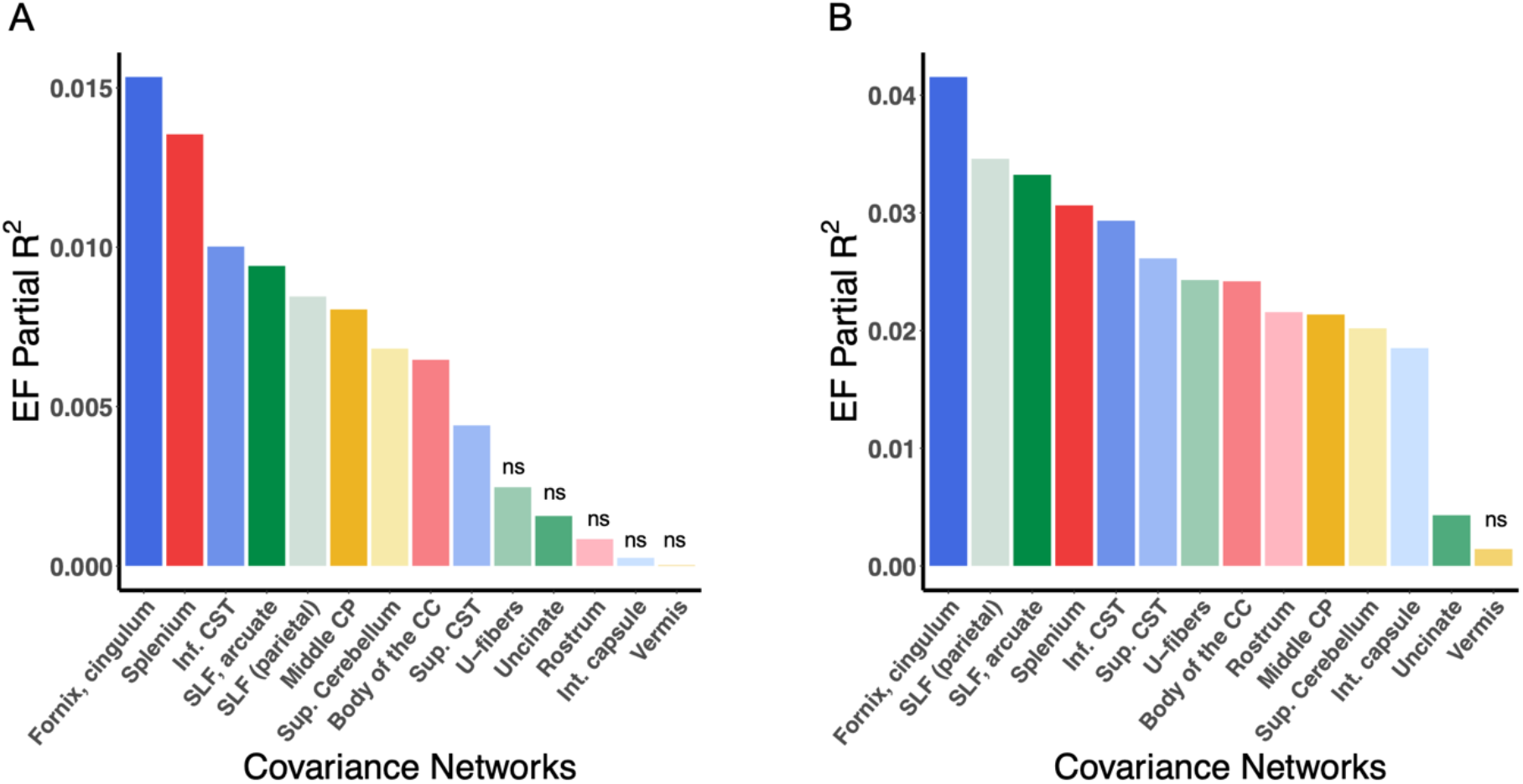
Effect sizes (partial *R*^*2*^) of executive function in each fiber covariance network controlling for total brain volume and maternal education. **A)** Bar graph depicting the effect size of executive function for each network (partial *R*^*2*^) while controlling for total brain volume. **B)** Bar graph depicting the effect size of executive function for each network (partial *R*^*2*^) while controlling for maternal education. Non-significant associations are marked by “ns”. Abbreviations: EF, executive function; CC, corpus callosum; SLF, superior longitudinal fasciculus; CST, cortico-spinal tract; Sup, superior; Int, internal; CP, cerebellar peduncle.

